# Whole genome scan reveals the multigenic basis of recent tidal marsh adaptation in a sparrow

**DOI:** 10.1101/360008

**Authors:** P. Deane-Coe, B. G. Butcher, R. Greenberg, I. J. Lovette

## Abstract

Natural selection acts on functional molecular variation to create local adaptation, the “good fit” we observe between an organism’s phenotype and its environment. Genomic comparisons of lineages in the earliest stages of adaptive divergence have high power to reveal genes under natural selection because molecular signatures of selection on functional loci are maximally detectable when overall genomic divergence is low. We conducted a scan for local adaptation genes in the North American swamp sparrow (*Melospiza georgiana*), a species that includes geographically connected populations that are differentially adapted to freshwater vs. brackish tidal marshes. The brackish tidal marsh form has rapidly evolved tolerance for salinity, a deeper bill, and darker plumage since colonizing coastal habitats within the last 15,000 years. Despite their phenotypic differences, background genomic divergence between these populations is very low, rendering signatures of natural selection associated with this recent coastal adaptation highly detectable. We recovered a multigenic snapshot of ecological selection via a whole genome scan that revealed robust signatures of selection at 31 genes with functional connections to bill shape, plumage melanism and salt tolerance. As in Darwin’s finches, BMP signaling appears responsible for changes in bill depth, a putative magic trait for ecological speciation. A signal of selection at BNC2, a melanocyte transcription factor responsible for human skin color saturation, implicates a shared genetic mechanism for sparrow plumage color and human skin tone. Genes for salinity tolerance constituted the majority of adaptive candidates identified in this genome scan (23/31) and included vasoconstriction hormones that can flexibly modify osmotic balance in tune with the tidal cycle by influencing both drinking behavior and kidney physiology. Other salt tolerance genes had potential pleiotropic effects on bill depth and melanism (6/31), offering a mechanistic explanation for why these traits have evolved together in coastal swamp sparrows, and in other organisms that have converged on the same “salt marsh syndrome”. As a set, these candidates capture the suite of physiological changes that coastal swamp sparrows have evolved in response to selection pressures exerted by a novel and challenging habitat.

## Introduction

Natural selection is one of the most fundamental evolutionary forces, solely responsible for generating present day adaptive diversity by sorting functional molecular variation within lineages to drive local adaptation. When selection is strong, and when adaptive phenotypic divergence influences reproductive isolation, locally adapted lineages may advance along the speciation continuum toward ecological speciation (Mayr 1963; Harrison 1991; Schluter 2000, 2001; Coyne and Orr 2004; Nosil et al. 2005; Rundle and Nosil 2005; Nosil 2012; Harrison 2012). This process can therefore be better understood by characterizing the molecular mechanisms underlying local adaptation. The functional effects of genes responsible for adaptive phenotypes offer one such window onto the machinery of evolution.

Recent studies have searched for genes under selection by conducting whole genome comparisons of closely related taxa (eg. Rubin et al. 2010; Cao et al. 2011; Jones et al. 2012; Malinsky et al. 2015; Burri et al. 2015; Lopes et al. 2016; Toews et al. 2016). Modern genome scans typically reveal heterogeneous patterns due to a combination of factors that include selection, gene flow, incomplete lineage sorting, and recombination rate variation (Payseur and Rieseberg 2014). Peaks that stand out from the genomic background are often considered “islands of divergence” (Turner et al. 2005; Seehausen et al. 2014). Because the most commonly used divergence metric (*F*_ST_) is a relative measure, sensitive to both allele frequency differences and within-lineage variation at a particular site, islands of divergence may represent impermeable regions in a semipermeable genome that have been shielded from gene flow by selection (Harrison 1986, 1990; Wu 2001; Turner et al. 2005; Kane et al. 2009) or regions that have been purged of molecular variation by selective sweeps (Cruickshank and Hahn 2014; Delmore et al. 2015). Both must be distinguished from background selection in low recombination regions, a confounding process also capable of generating *F*_ST_ peaks (Charlesworth et al. 1993; Charlesworth 1998; Noor and Bennet 2010; Nachman and Payseur 2012; Cruickshank and Hahn 2014; Burri et al. 2015; Van Doren et al. 2017).

Populations in the earliest stages of ecological divergence across an environmental gradient provide an opportunity to characterize local adaptation genes (Harrison and Larson 2014; Seehausen et al. 2014). Recent divergence times and ongoing gene flow make adaptive regions of the genome maximally detectable against a highly similar genomic background (Seehausen et al. 2014; Payseur and Rieseberg 2016). The North American swamp sparrow (*Melospiza georgiana*) fits these criteria well, making it a tractable system in which to identify molecular targets of selection during adaptive divergence. Multiple lines of evidence suggest that swamp sparrows have recently expanded their range to colonize brackish coastal tidal marshes, as much of the northeastern and mid-atlantic coast was under an ice sheet at the Last Glacial Maximum. Mid-atlantic coasts were also hydrologically unstable during much of the Pleistocene due to glacial outflow from rivers and streams (Malamud-Roam et al. 2006), and tidal marshes require sediment accretion from gradually rising sea levels in order to establish (Pethick 1984; Warren and Niering 1993). Coastal swamp sparrows have therefore likely colonized this novel habitat within the last 15,000 years (Greenberg and Droege 1990). In the present, habitat isolation is the only potential barrier to gene flow between coastal and inland swamp sparrows, and they continue to interbreed at the ecotone between inland freshwater and coastal brackish habitats (Greenberg et al. 2016; Deane-Coe et al. 2018).

### Local adaptation to tidal marshes

Despite a very recent divergence time and active contemporary gene flow with inland populations, coastal swamp sparrows are morphologically, ecologically and behaviorally distinct from inland populations (Greenberg and Droege 1990; Deane-Coe et al. 2018). At least 10 well-characterized traits distinguish coastal swamp sparrows that breed in tidal marshes (*M. g. nigrescens*) from the more common inland freshwater subspecies (*M. g. georgiana*), including a tolerance for salinity, a deeper bill, and more melanic plumage. (Figure 1; Greenberg and Droege 1990; Grenier and Greenberg 2005; Olsen et al. 2013). Common garden experiments have revealed that differences in bill depth and plumage melanism between inland and coastal swamp sparrows persist when they are reared together in the laboratory, indicating that these components of phenotypic divergence have a heritable genetic basis (Ballentine and Greenberg 2010), and the adaptive value of these traits have been explicitly tested and demonstrated in natural populations (summarized in Deane-Coe et al. 2018). Physiological and molecular mechanisms responsible for salt tolerance, bill depth and melanism have been characterized in many other species, providing a robust functional genomic literature from which to extract candidate genes, pathways and processes that may underlie these traits in swamp sparrows.

**Figure 1.**
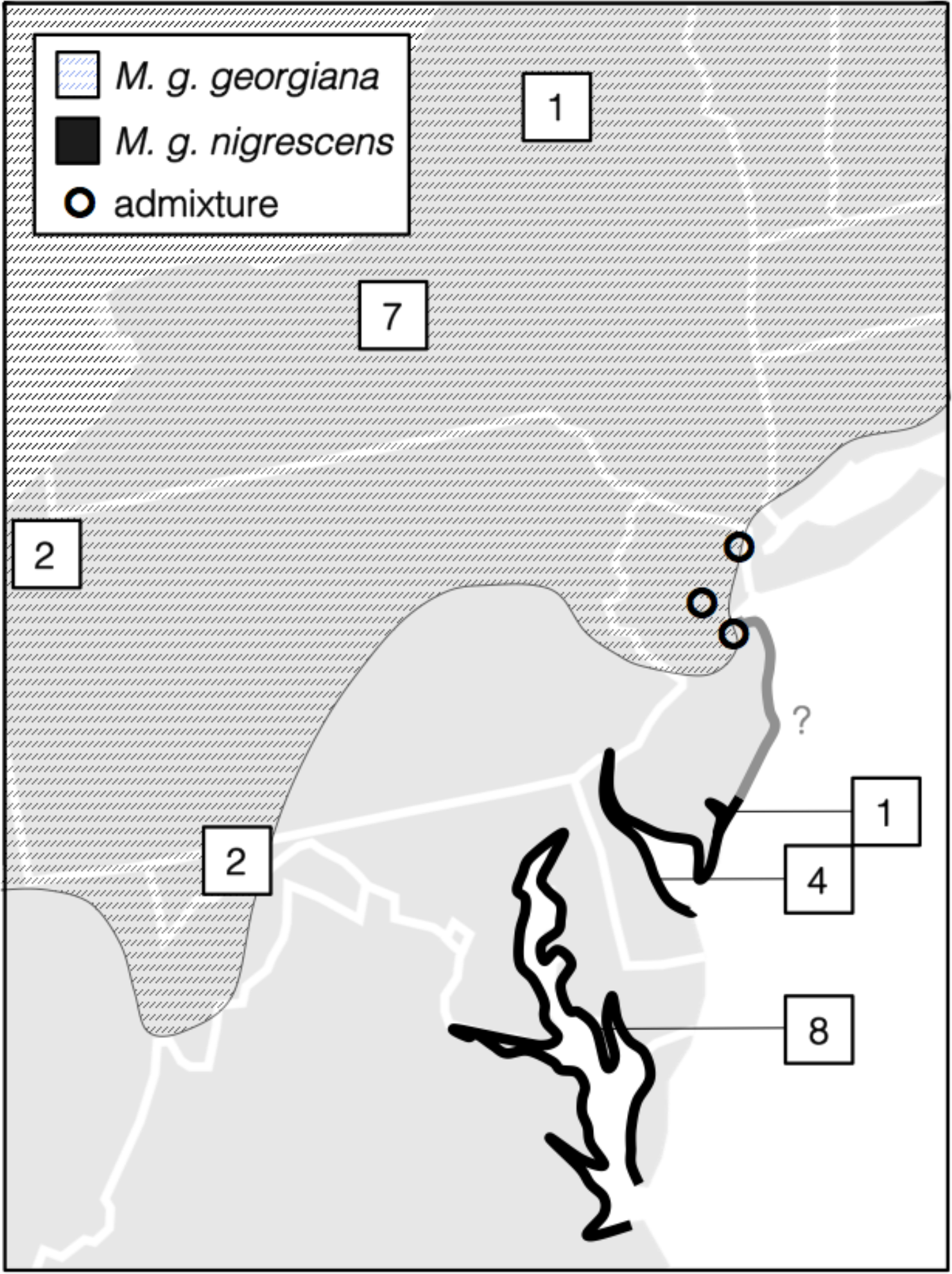
Breeding range of inland and coastal swamp sparrow subspecies in the mid-atlantic US, and their contact zone at the ecotone between freshwater and brackish tidal marshes in northern New Jersey (NJ). Tidal marsh habitat along the NJ shore contains previously unsampled (and therefore as-yet undiagnosed) populations (grey line). Numbers indicate samples from each site.

### Salinity tolerance

Tidal salinity poses a fluctuating osmoregulatory challenge that all organisms breeding in tidal marshes must contend with physiologically, and it is one of the reasons that tidal marsh habitats are predominantly inhabited by specialist taxa (Correll et al. 2016). As tidal marsh specialists, coastal swamp sparrows are somehow able to tolerate increased salt intake from brackish drinking water and from foraging for invertebrate prey in brackish habitats (Greenberg et al. 2006). A coastal individual’s exposure to dietary salinity, and the degree of osmoregulatory stress they must tolerate, is a product of where they drink water and where their preferred invertebrate prey live (Figure 2; Goldstein 2006).

**Figure 2.**
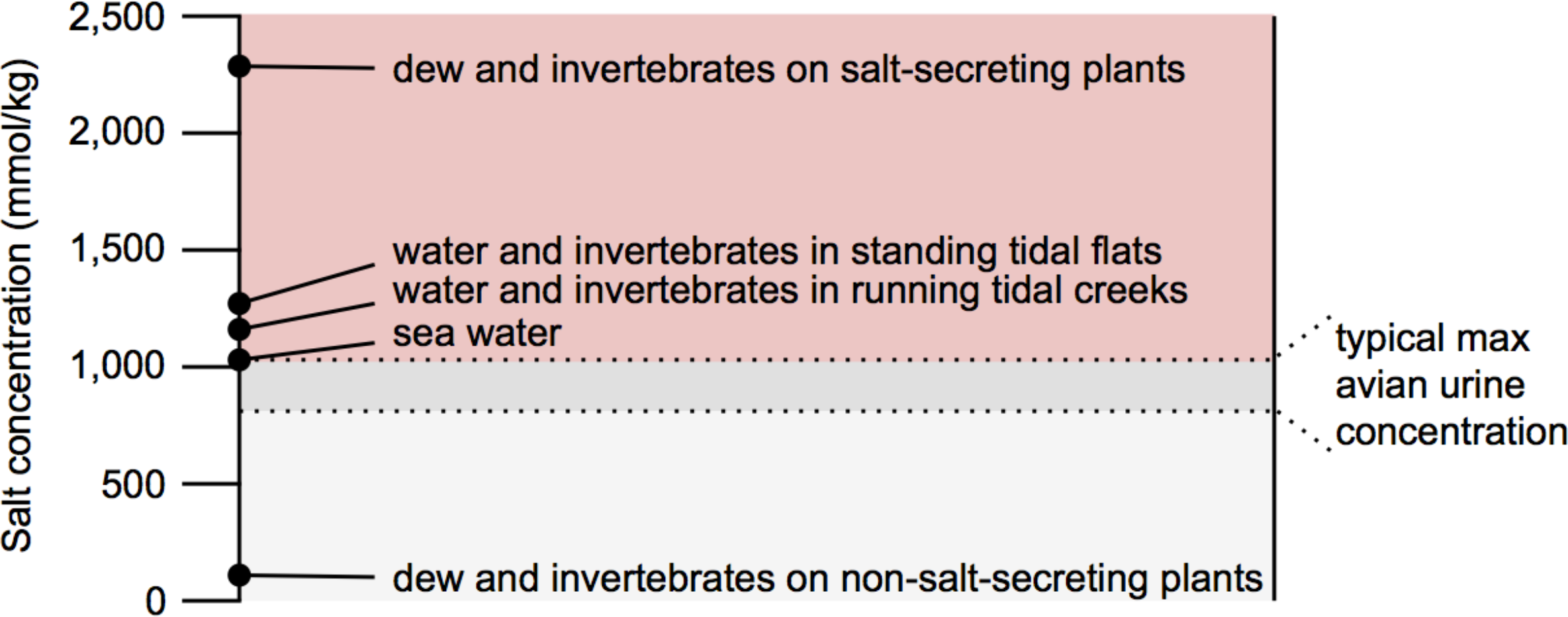
Comparison of salt concentration in different food and water sources in a tidal marsh with the maximum salt-concentrating ability of the avian kidney (copied from Goldstein et al. 1990, and adapted to include information from Goldstein 2006). Tidal marsh sparrows must evolve specialized behavioral and/or physiological mechanisms of salt tolerance, since most available food or water sources contain salt concentrations that exceed the maximum concentration of typical avian urine.

Pelagic seabirds have specialized glands to concentrate and excrete salt (Heatwole and Taylor 1987), as does the saltmarsh-breeding clapper rail (Olson 1997), but passerine birds like sparrows have no such glands. Instead, passerines in saline habitats must tolerate salt through behavioral modifications that limit salt ingestion and physiological modifications that eliminate excess dietary salt via the kidney and intestine (Goldstein 2006). In a controlled comparison in a laboratory setting, sparrow species that breed in saltmarshes decreased drinking rate and maintained body mass better at higher salinity compared to inland sparrows (Bartholemew and Cade 1963). This decrease in drinking rate demonstrates that saltmarsh-adapted sparrows possess behavior-modifying mechanisms that aid osmoregulation, and their ability to maintain body mass demonstrates that they have physiological mechanisms to prevent water loss. For example, anatomical comparisons revealed that saltmarsh-adapted sparrows have structural adaptations to reduce salt reabsorption by the intestine (a smoother colon with a reduced brush border; Goldstein et al. 1990), and enlarged kidneys with an enhanced system for urine concentration (more medullary cones and loops of Henle in the nephron; Goldstein 2006).

The genetic changes responsible for these specific behavioral and physiological adaptations in saltmarsh-adapted sparrows are not yet known. To date, molecular mechanisms of salt tolerance have largely been characterized for species that cannot behaviorally escape salinity, such as plants (eg. Shi et al. 2000, Xue et al. 2004; Kane et al. 2007), fish (eg. Hyndman and Evans 2007, 2009; Rengmark et al. 2007; Purcell et al. 2008) and aquatic invertebrates (eg. Patrick et al. 2000). In saline environments these organisms may increase the transport and excretion of Na^+^ and Cl^-^ to maintain optimal osmotic balance. In plants and fish, this is often achieved by increasing the expression, distribution or affinity of Na^+^ and Cl^-^ ion channels (Munns 2005; Hauser and Horie 2010). Much of this regulation occurs in the distal convoluted tubule (DCT) of the kidney, where the final excreted concentration of ions is determined via transporter-mediated reabsorption (de Baaij et al. 2015). In plants, ion channels expressed in the membranes of intracellular vesicles sequester and compartmentalize excess salt under salt stress conditions (Batelli et al. 2007). Pathways that regulate vesicle trafficking and turnover can confer salt tolerance by accelerating degradation or secretion of products sequestered in vacuoles, endosomes or lysosomes (Mazel et al. 2004). Organisms must also reduce intestinal permeability and transport of other ions like Mg^2+^ to maintain homeostasis without wasting water during excretion (Grosell et al. 2011; Esbaugh et al. 2014; Romani et al. 2007). Finally, in environments like tidal marshes where salinity stress changes throughout the tide cycle, rates of ion transport may be temporarily modified via vasoconstriction or vasodilation of the renal arteries or glomerular capillaries (Hyndman 2015).

### Deep bills as radiators

For coastal birds, deeper bills offer a larger surface area for convective radiative heat loss, reducing thermal stress in the sun-exposed, water-limited conditions that characterize tidal marshes (Greenberg et al. 2012). Natural selection for deeper bills in coastal swamp sparrows has, in turn, driven divergence in song (Ballentine 2006): coastal males are biomechanically constrained from singing the broad-bandwidth trills that inland females prefer, but coastal females prefer the alternative songs of coastal males, leading to non-random mating (Ballentine et al. 2013a, 2013b). This mirrors the evolution of incipient reproductive isolation in Darwin’s finches and crossbills, where ecological selection favored larger bills in some populations, which in turn exerted constraints on song and promoted non-random mating (Podos 2001; Podos et al. 2004; Huber and Podos 2006; Huber et al. 2007; Smith and Benkman 2007). Swamp sparrow bill depth may therefore act as a “magic trait”: a mechanism by which natural selection itself drives ecological speciation between inland and coastal lineages over time (Gavrilets 2004).

Genetic determinants of bill depth variation in birds are already well understood because this trait plays a role in several evolutionarily and agriculturally important model systems. Bone morphogenetic proteins (BMPs) are members of the Transforming Growth Factor β (TGFβ) superfamily, and BMP signaling is responsible for variation in bill depth in Darwin’s finches, chickens, ducks and quail via their action during craniofacial development (Abzhanov et al. 2004; Wu et al. 2004, 2006; Brugmann et al. 2010). Over-expression of the growth factor BMP4 in the prenasal cartilage of developing chicken embryos caused those individuals to grow larger, deeper bills (Abzhanov et al. 2004; Wu et al. 2004). Other genes acting during development can lead to craniofacial deformities in humans, and several of these have also been implicated in bill shape evolution in birds (eg. Lamichhaney et al. 2015).

### Increased melanism for feather preservation and territoriality

Coastal swamp sparrows may have evolved more melanic plumage than their inland relatives because feather-degrading bacteria are more abundant in in humid coastal habitats like tidal marshes, and feathers with a higher concentration of melanins resist bacterial degradation better (Peele et al. 2009). Increased melanism also enhances the size and conspicuousness of their black forehead patch, a plumage badge that indicates dominance and aids males in territorial competition. Male-male competition is more intense for coastal swamp sparrows in comparison to inland swamp sparrows because breeding densities are very high in tidal marshes (Olsen et al. 2008a). Although it has been observed that anoxic iron sulfides cause tidal marsh sediments to be darker on average than inland marshes, inspiring speculation that melanism may also enhance crypsis (Greenberg and Droege 1990), this hypothesis has yet to be tested in swamp sparrows.

Melanic plumage is generated via the action of genes in the melanogenesis pathway (Mundy 2005), and via the function of specialized cells in the skin called melanocytes. Melanocytes contain melanosomes, specialized vesicles that collect the products of melanogenesis, melanins, and deploy them during feather development (Yu et al. 2004). Melanocyte function has been heavily studied in the context of feather patterning (eg. Lin et al. 2013), and several different members of the melanogenesis pathway have been implicated as functional candidates for melanic plumage or pelage in several different species. For example, mutations in the melanocortin-1 receptor (MC1R; Theron et al. 2001; Uy et al. 2016; reviewed in Mundy 2005) or it’s antagonist, agouti signalling protein (ASIP; Manceau et al. 2011; Poelstra et al. 2014; Toews et al. 2016; Campagna et al. 2016), appear responsible for the switch between phaeomelanin (brown) and eumelanin (black) production in a range of different birds and mammals (Hubbard et al. 2010).

### Melanic plumage and large bills, a saltmarsh “syndrome”

Coastal subspecies of the swamp sparrow, saltmarsh sparrow, seaside sparrow, song sparrow and marsh wren are all more melanic than their inland relatives (Grinnell 1913; Von Bloeker 1932; Phillips 1986; Greenberg and Droege 1990; Luttrell et al. 2014). Saltmarsh melanism has also been described for black rails, shrews, voles, harvest mice, and the gulf saltmarsh snake (Von Bloeker 1932; Neill 1958; Pettus 1963; Conant and Lazell 1973; Myers 1988; Gaul 1996). In addition to this pervasive pattern of melanism in saltmarsh taxa, all 10 North American sparrow species or subspecies that breed in salt marshes have evolved larger bills than their closest inland relative (Grenier and Greenberg 2005). For example, in savannah sparrows (*Passerculus sandwichensis*), the tidal marsh subspecies *rostratus* has a much deeper bill than inland savannah sparrows (Grenier and Greenberg 2005) and in song sparrows (*Melospiza melodia*) the tidal marsh subspecies *maxillaris* has both a larger bill and more melanic plumage than inland song sparrows (Marshall 1948; Grenier and Greenberg 2005; Luttrell et al. 2014). Increased summer temperatures in tidal marshes relative to inland marshes offers an adaptationist explanation for this repeated evolution of large-billed forms, because increased bill volume improves radiative heat loss (Greenberg et al. 2012). Extensive phenotypic convergence would seem to constitute strong evidence for the adaptive value of deep bills and melanic plumage for a wide range of organisms living in saltmarsh habitats. As a recently diverged, tractable system that offers high detectability for adaptive variation in the genome, the swamp sparrow provides a means to understand a very taxonomically broad pattern of adaptive diversity.

We conducted a scan to identify peaks of divergence in the genomes of inland and coastal swamp sparrows, and to determine whether peaks coincide with the locations of candidate genes. We defined a candidate gene as any gene with one or more functional connections to previously known physiological, cellular or molecular mechanisms of salinity tolerance, bill morphogenesis or melanogenesis (Table 1). Where possible, we differentiate between peaks in several different locations relative to transcription start sites of candidate genes, representing functionally distinct categories of molecular evolution: 1) in coding regions, 2) immediately upstream at the putative promoter or primary enhancer, 3) in introns of the target gene or neighboring genes, or 4) in adjacent non-coding regions. The last two categories represent possible sites of secondary regulatory elements (eg. “shadow enhancers”; Hong et al. 2008), which may influence the expression of target genes many kilobases away (eg. Bishop et al. 2000; Lettice et al. 2002).

**Table 1.**
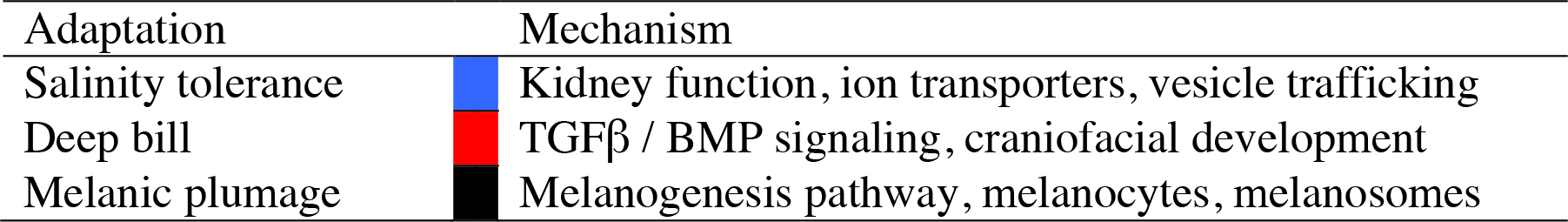
Candidate mechanisms for three main categories of coastal adaptation.

| Adaptation | Mechanism |
| --- | --- |
| Salinity tolerance | Kidney function, ion transporters, vesicle trafficking |
| Deep bill | TGFβ / BMP signaling, craniofacial development |
| Melanic plumage | Melanogenesis pathway, melanocytes, melanosomes |

## Methods

### Reference genome construction

We sequenced the genome of a male swamp sparrow in breeding condition, sampled 14 June 2014 from an active territory at a coastal breeding site in Fishing Bay Wildlife Management Area (MD) on the eastern shore of the Chesapeake Bay (38°24’36”N 76°00’09”W). This site represents the most southern coastal breeding population sampled to date, located at the greatest distance from the southern range edge of the inland subspecies. The skin specimen is vouchered at the Cornell University Museum of Vertebrates (Accession no. N). We extracted 5.5 ug of DNA from this sample using the DNeasy kit (Qiagen) and submitted it to the Genomics and Epigenomics Core Facility of the Weill Cornell Medicine Core Laboratories Center (CLC) for whole genome library preparation and sequencing. Weill CLC prepared two mate-pair libraries with 3kb and 8kb inserts using Nextera Mate Pair library preparation kits, and one 180bp fragment library using a Nextera DNA sample preparation kit (Illumina, San Diego, CA, USA) (Table S1). We confirmed target fragment size and concentration for each library with a Bioanalyzer (Agilent, Santa Clara, CA, USA), and paired end sequencing (2×100bp) was performed on the Illumina HiSeq2500 platform. Both large-insert mate pair libraries were sequenced on a single lane, and the fragment library was sequenced across two lanes.

We filtered reads failing the Illumina chastity filter and assembled remaining reads (3kb 507,715,832 reads; 8kb 466,743,564 reads; fragment 542,406,170 reads) into scaffolds using the ALLPATHS-LG pipeline, which additionally incorporates read quality and base pair uncertainty into the assembly algorithm (Gnerre et al. 2011; Ribeiro et al. 2012). We also annotated the genome using MAKER 2.32 (Cantarel et al. 2008), creating gene models from Ensembl protein and cDNA databases for the zebra finch (*Taeniopygia guttata*) and gene predictions from the program SNAP (Korf 2004).

### Genome resequencing and genotyping

We compiled 24 blood or tissue samples from breeding males at allopatric coastal and inland sites (N=12 per subspecies; Figure 12; Table S1). Previous work has demonstrated an absence of genetic substructure within either subspecies, and no effect of isolation by distance at this geographic scale (Deane-Coe et al. 2018). We documented phenotypes by taking a complete set of high resolution photographs for all birds banded in 2012-2014, using a Canon DSLR camera against a standardized color reference target (Xrite Digital SG Colorchecker), and collected standard morphological data for all banded birds. Blood was preserved in Queen’s lysis buffer, tissue was preserved in ethanol, and both were stored at -20°C. We extracted DNA from all samples using the DNeasy kit (Qiagen), quantified concentrations on a Qubit (Invitrogen, Carlsbad, CA, USA), and balanced concentrations to 2.3 ng/ul (119 ng total in 52.5 ul) across samples using either dilution or vacuum filtration.

We prepared a whole genome library in 350bp sonicated fragments for each individual using the Illumina TruSeq Nano library preparation kit, and performed paired-end sequencing of all 24 libraries on one lane of an Illumina NextSeq500 platform (2×150bp). One coastal sample (SWSP12, Egg Harbor NJ) did not result in high quality reads and was dropped prior to read processing and genotyping. We used the program AdapterRemoval (Lindgreen 2012) to trim Ns from reads, truncate read pairs that contain adapter sequence, and collapse any overlapping read pairs into a single read. We required that every read have a Phred quality score and a minimum length of at least 20.

Bowtie2, a gapped-read aligner (Langmead and Salzberg 2012), was used to map paired reads and unpaired collapsed reads from each individual to the reference assembly using the “very sensitive local” flag, which applies the following parameters to the alignment algorithm: a maximum effort of 20 seed extension attempts and 3 re-seed attempts, 0 mismatches allowed, a seed substring length of 20, and the interval between seed substrings defined by a square root function with a constant of 1 and coefficient of 0.5. We used Samtools (Li et al. 2009) to convert individual alignment files from sequence alignment/map format (SAM) to the binary version (BAM), and to index the fasta file for the reference genome. We applied tools from the program Picard (Wysoker et al. 2012) to add sequencing group information to individual BAM files prior to genotyping, and coordinate contig names across the dataset. We flagged individual fragments that had been sequenced more than one time as duplicates, so that they did not mislead downstream genotyping, and indexed the resulting flagged alignments. We also created a dictionary of contig names and sizes for the reference genome.

Individual genotypes and genotype likelihoods were assigned using the UnifiedGenotyper from Genome Analysis Toolkit (GATK; DePristo et al. 2011). When identifying variant sites and assigning genotypes we required a minimum Phred-scaled base quality score of 17, and a minimum confidence threshold of 10 (34,358,667 variant sites from the alignments met these criteria). We then used VCFtools (Danecek et al. 2011) to apply more stringent filtering, requiring that each SNP be genotyped for all samples, have a minimum mapping quality score of 20, and a minimum mean depth of 5 reads across all individuals (13,819,090 SNPs).

### Quantifying diversity, differentiation, and divergence

For each SNP we estimated Weir and Cockerham *F*_ST_ and nucleotide diversity (π) using VCFtools. We calculated 50 kb sliding window estimates of *F*_ST_ and π using a window step size of 10 kb, and plotted them using the Manhattan plot R package qqman (Turner 2014) to scan for regions of the genome that differed from the mean genomic background value in these stats. In comparison to blocked sliding windows (in which each regional estimate is independent), sliding windows with overlapping steps provide smoothed estimates that reduce noise in the dataset while accurately capturing any signal of gradual change across regions of interest. We defined differentiated regions as 50 kb windows for which mean *F*_ST_ was more than five standard deviations above the genome-wide mean, considering only the largest and best quality scaffolds in our assembly (scaffolds 0 to 150). We also scanned for long runs of homozygosity (LROH) in each differentiated region with VCFtools to test for the presence of a recent selective sweep, and calculated Tajima’s D for 1 kb windows across the scaffold to infer other signatures of selection or demography from allele frequency spectra in each window. Positive values of Tajima’s D can result from excess polymorphism at intermediate frequencies, a pattern that can be generated at individual loci via balancing selection or genetic structure within the sampled population. To distinguish among these alternatives we tested for population structure by estimating *F*_ST_ between coastal sites using the SNPs that defined each peak with positive Tajima’s D. When sites under peaks were invariant within coastal swamp sparrows, we expanded the region used to estimate *F*_ST_ by 1Mb (500kb in either direction). Negative values of Tajima’s D indicate an abundance of rare alleles, a pattern that can be generated at individual loci via mutational accumulation of variation following a selective sweep, or genome-wide via population expansion after a bottleneck (Tajima 1989; Fu and Li 1993; Jensen et al. 2005).

### Functional annotation

We mapped the swamp sparrow sequence under each *F*_ST_ peak to the zebra finch (*Taeniopygia guttata*) genome assembly (Taeniopygia_guttata-3.2.4, reference Annotation Release 103) using BLASTn (NCBI). We required that sequences mapped well (>70% query coverage, >80% sequence identity, E value 0.0) to a single location in the assembly in order for the genes under that peak to be considered candidates. Assuming conservation of synteny with the zebra finch, we queried NCBI for the location of the centromere of each chromosome harboring a divergence peak in swamp sparrows to assess whether peaks coincided (<1MB) with these regions of low recombination. We referred to gene entries from Entrez, UniProtKB/Swiss-Prot, GeneCards and Tocris databases to assign functional descriptions to each candidate gene located under or adjacent to the peak.

## Results

### Coastal swamp sparrow reference genome

Our ALLPATHS-LG reference assembly contained 4,778 scaffolds, with an N50 scaffold size of approximately 10Mb and an estimated genome size of 1.2Gb (Table S2; Table S3).

### Regions of high divergence

Genome-wide divergence between inland and coastal swamp sparrows was very low (mean *F*_ST_=0.015; Figure 3). Average genome-wide estimates of Tajima’s D were negative in both inland (*D* = -0.73) and coastal birds (*D* = -0.42), representing a genome-wide signal of population expansion after a recent bottleneck in both populations. Restricting our scan to the largest scaffolds in our assembly (scaffolds 0-150), we identified 41 regions where the 50kb sliding window estimate of *F*_ST_ exceeded five standard deviations above the mean (0.08; Figure 3). The average size of a divergence peak was 171kb (± 294kb), the largest being 1.5Mb (scaffold 148). These windows contained highly differentiated SNPs (max *F*_ST_=0.8), and deviations from the genomic baseline in π and Tajima’s D. BLAST mapping of the sequence underlying each peak revealed that, out of the 138 genes located under peaks, 15 (11%) had functional effects that made them candidates for coastal adaptation. An additional 16 candidates were located adjacent to peaks. In total, 31 genes with functional connections to salt tolerance, bill depth or melanogenesis were located immediately under or adjacent to 22 of the 41 peaks characterized genome-wide (Figures 4-7; Table 2; Table S4; Figures A2.2-2.4). The remaining 19 peaks (Table S4) either did not contain candidate genes for coastal adaptation (N=16), or did not BLAST well to an annotated location in the zebra finch assembly (N=3).

**Figure 3.**
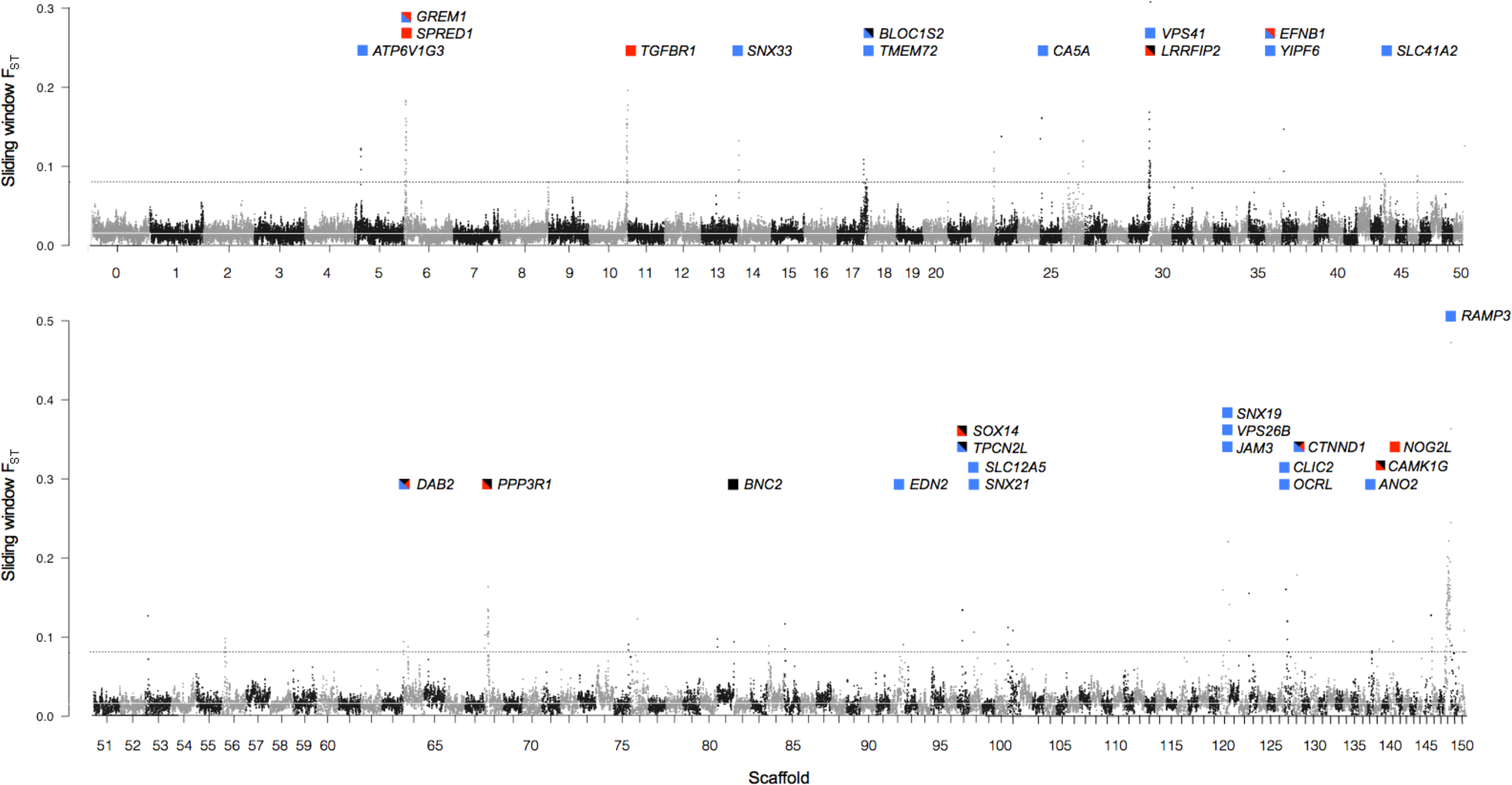
Genome-wide Manhattan plot between inland and coastal swamp sparrows, based on 50 kb sliding window estimates of *F*_ST_ across the largest scaffolds in our genome assembly (0-150). Genome-wide average *F*_ST_ is very low (*F*_ST_=0.015; white line). Colored flags represent the location and functional annotation of candidate genes for coastal adaptation located at *F*_ST_ peaks more than five standard deviations (0.013) above the mean (outlier threshold=0.08; dashed line) (Table 4). Refer to Table 1 for color key.

**Figure 4.**
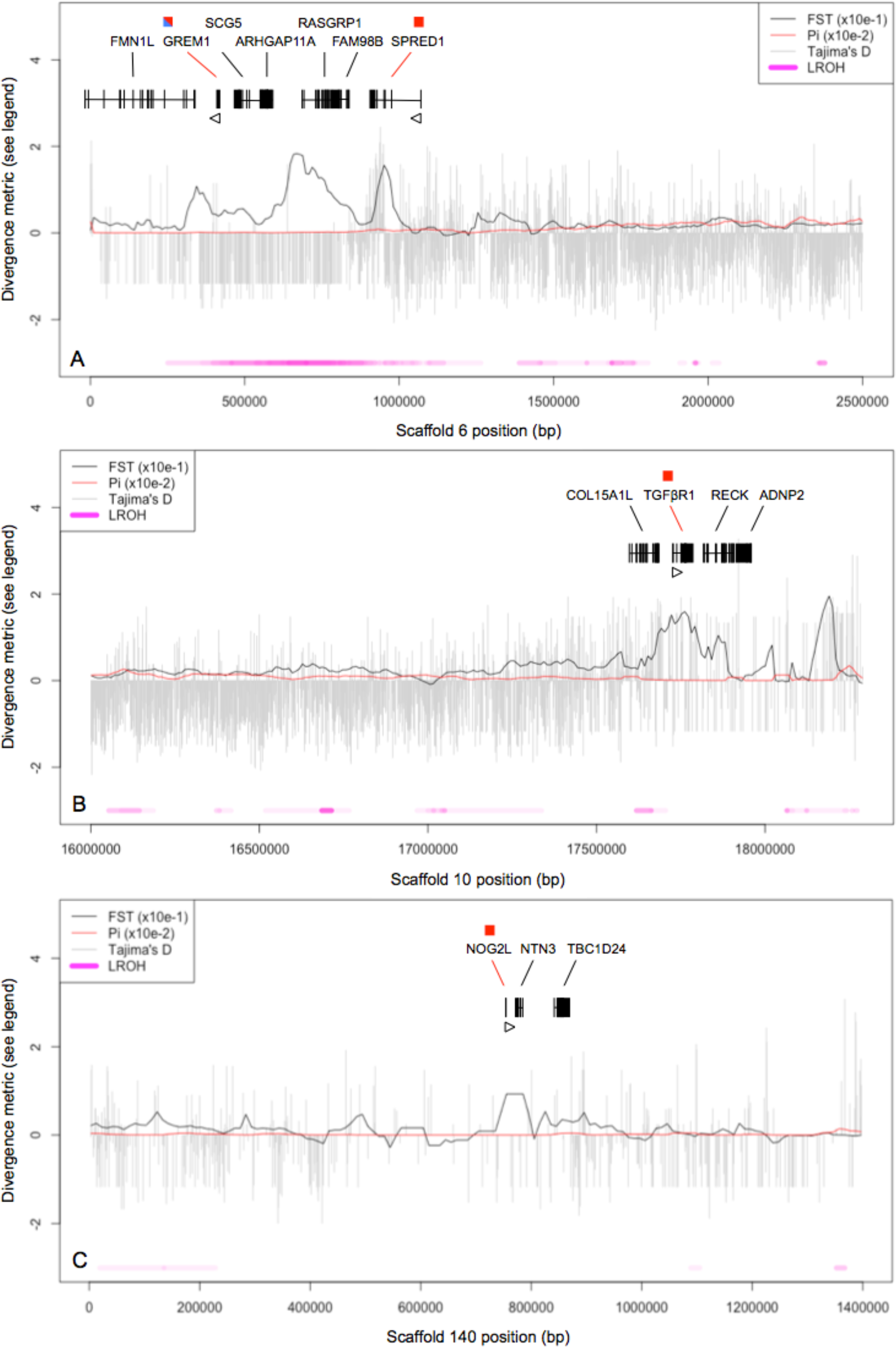
*F*_ST_ peaks on scaffolds 6 (A), 10 (B) and 140 (C) coincide with regions of low or zero nucleotide diversity (π) in coastal and inland swamp sparrows, consistent with recent selective sweeps, and contain three members of the BMP signaling cascade. On scaffold 6 (A), elevated F_ST_ also coincides with long runs of homozygosity (LROH) and an excess of rare alleles (Tajima’s D < 0) in coastal swamp sparrows, further diagnostic of a sweep. Sites lacking Tajima’s D estimates are invariant in coastal swamp sparrows. For LROH, color intensity corresponds to the count of coastal individuals with runs of homozygosity at that position.

**Figure 5.**
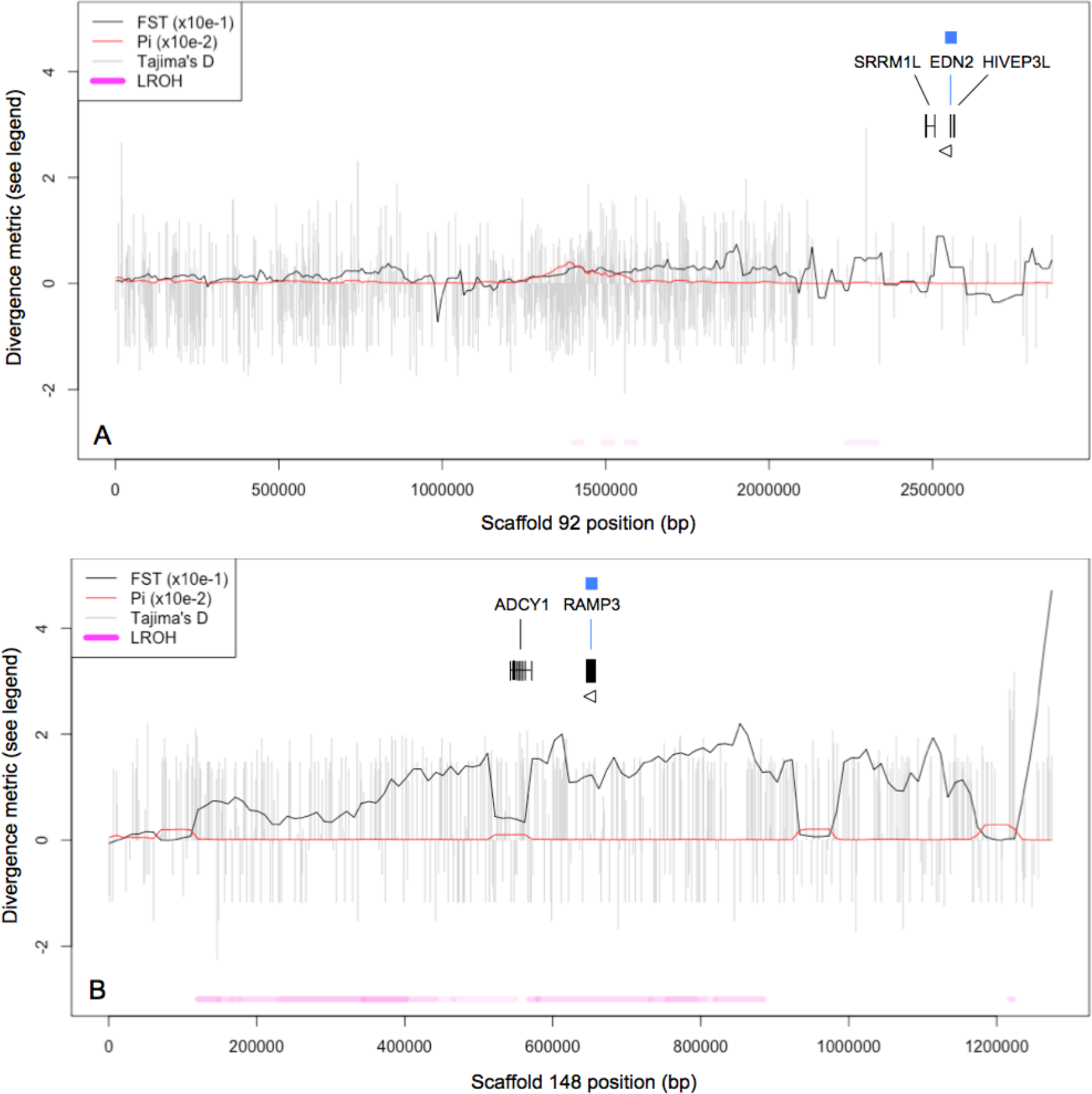
*F*_ST_ peaks on scaffolds 92 (A) and 148 (B) map to regions containing genes that encode kidney vasoconstriction and dilation hormones. Both show reduced nucleotide diversity (π). For LROH, color intensity corresponds to the count of coastal individuals with runs of homozygosity at that position.

**Figure 6.**
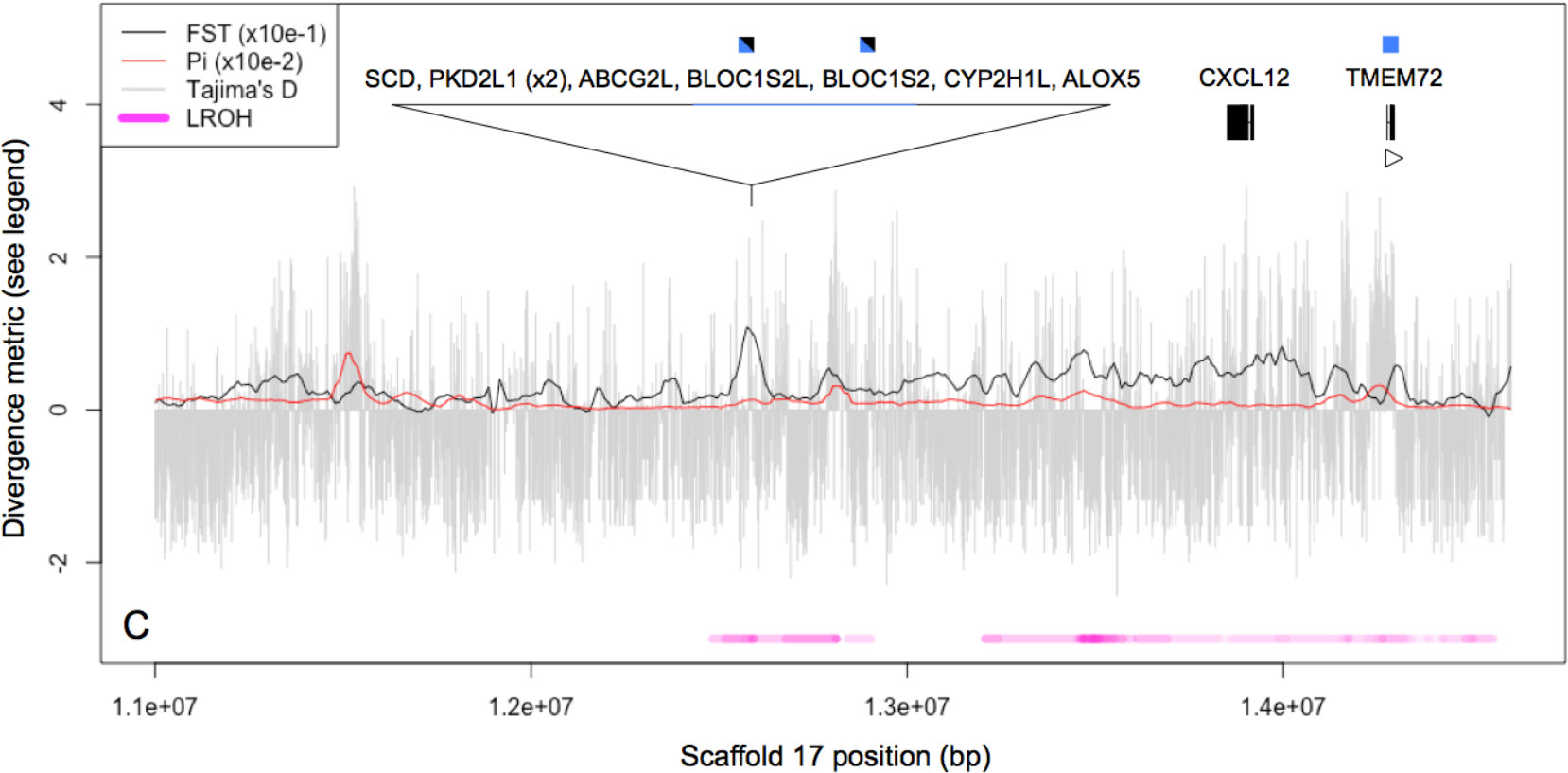
*F*_ST_ peaks on scaffolds 17 map to regions containing salinity tolerance genes that also show runs of homozygosity (LROH) and divergence in Tajima’s D relative to the genomic background. For LROH, color intensity corresponds to the count of coastal individuals with runs of homozygosity at that position. The candidate BLOC1S2 may exert pleiotropic effects on both salinity tolerance and melanogenesis.

**Figure 7.**
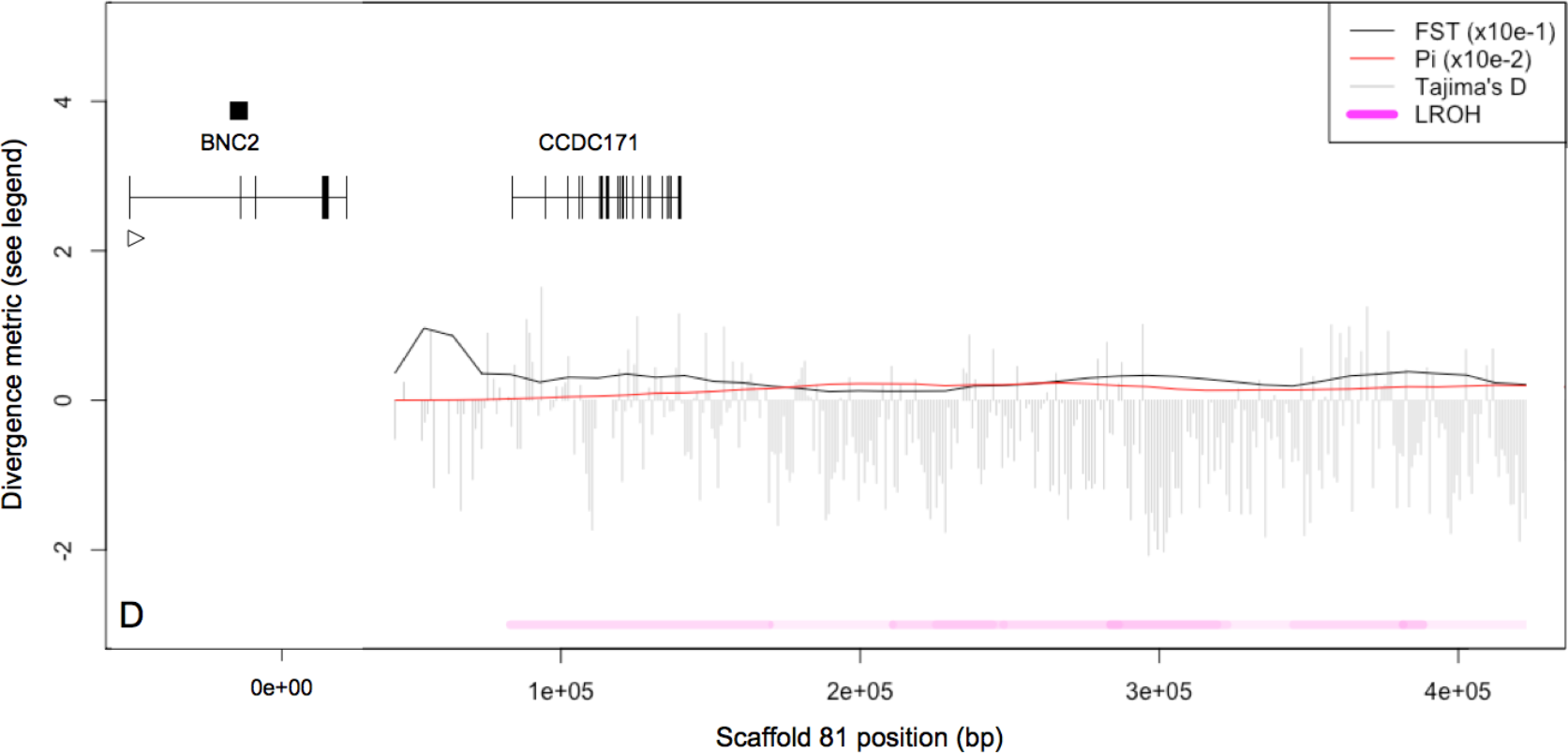
The *F*_ST_ peak on scaffold 81 maps to a region containing a candidate gene for melanic plumage that influences skin color saturation in human populations. Runs of homozygosity (LROH) are present across this small scaffold in several coastal individuals. For LROH, color intensity corresponds to the count of coastal individuals with runs of homozygosity at that position.

Most candidate genes were located in regions with very low or zero π relative to the genomic background (25/31; Figure 4-2.5; Table 2; Figures A2.1-A2.4). A different but overlapping subset of candidates (13/31) coincided with runs of homozygosity in coastal swamp sparrows. All 31 *F*_ST_ peaks exhibited concomitant runs of homozygosity, reduced nucleotide diversity, or both. Positive estimates of Tajima’s D were explained by allele frequency divergence between coastal breeding sites for three of nine loci, but the remainder exhibited no structure within coastal swamp sparrows (*F*_ST_ < 0.1; Table S5). Most peaks containing candidate genes were not located in putative centromeric regions with low recombination rates (25/31).

**Table 2.** Candidate genes for coastal adaptation located at *F*_ST_ peaks in our genome scan (N=31). Regions containing these genes exhibited runs of homozygosity (H), reduced nucleotide diversity (π), or both. Negative Tajima’s D is consistent with a sweep, and positive D indicates balancing selection or structure among coastal sites at that locus. Signatures of selection at coding regions (EXON), proximal regulatory elements like primary enhancers (1°RE) or downstream/distal elements like secondary enhancers (2°RE) are given for peaks with sufficient resolution.

| Gene | Function | H | $\pi$ | D | Peak location |
| --- | --- | --- | --- | --- | --- |
| *SLC41A2* | Ion channel: plasma membrane $Mg^{2+}$ transporter | Y | ↓ | + | |
| *SLC12A5* | Ion channel: neuron-specific $Cl^-$ transporter | | ↓ | | |
| *CLIC2* | Ion channel: membrane-inserting $Cl^-$ transporter | Y | ↓ | + | |
| *ANO2* | Ion channel: $Ca^{2+}$ activated $Cl^-$ channel | Y | ↓ | + | 1°RE |
| *TMEM72* | Putative ion channel: expressed in kidney where $Na^+$ reabsorbed | Y | | ± | 2°RE |
| *TPCN2L* | Ion channel: expressed in kidney, affects human hair color |  | ↓ |  | 2°RE |
| *ATP6V1G3* | Vesicle trafficking: catalytic subunit of a vesicular ATPase $H^+$ pump | Y | | ± | 2°RE |
| *VPS41* | Vesicle trafficking: regulator of vesicle transport to lysosomes | Y |  | - | EXON |
| *VPS26B* | Vesicle trafficking: regulator of vesicle cargo recycling |  | ↓ |  | 1°RE |
| *SNX19* | Vesicle trafficking: regulator of vesicle trafficking and endocytosis |  | ↓ |  | 2°RE |
| *SNX21* | Vesicle trafficking: regulator of vesicle trafficking and endocytosis |  | ↓ |  |  |
| *SNX33* | Vesicle trafficking: regulator of vesicle trafficking and endocytosis |  | ↓ |  |  |
| *YIPF6* | Vesicle trafficking: associated with vesicle budding and transport | ~ | ↓ |  |  |
| *OCRL* | Vesicle trafficking: regulator of vesicle, lysosome, endosome traffic |  | ↓ |  |  |
| *BLOC1S2* | Vesicle trafficking: produces lysosomes and melanosomes | Y | ~ | ± |  |
| *DAB2* | Vesicle trafficking: regulator of vesicle traffic, CFTR, TGFβ/WNT signal | Y | | - | |
| *EDN2* | Kidney nephron vasoconstriction: the hormone endothelin 2 |  | ↓ |  | 2°RE |
| *RAMP3* | Kidney nephron vasodilation: a regulator of the hormone adrenomedullin | Y | ↓ | + | 1°RE / 2°RE |
| *CA5A* | Excretion: modifies water balance during excretion, affects ureagenesis |  | ↓ |  |  |
| *JAM3* | Solute permeability: adhesion molecule, prevents intracellular solute flow |  | ↓ |  | 1°RE |
| *CTNND1* | Solute permeability: assoc. with cadherin, affects adhesion, WNT signal |  | ↓ | - |  |
| *GREM1* | TGFβ signaling: BMP4 binding antagonist, affects bill/kidney development | Y | ↓ | - | 2°RE |
| *NOG2L* | TGFβ signaling: BMP4 binding antagonist | | ↓ | | 2°RE |
| *TGFBR1* | TGFβ signaling: receptor for TGFβ ligands, phosphorylates SMAD2 | Y | ↓ | + | EXON |
| *SPRED1* | FGF signaling: antagonist of fibroblast growth factors, affects face shape | Y | ↓ | ± | EXON |
| *LRRFIP2* | WNT signaling: may help activate WNT/β-catenin signaling | Y | | - | |
| *PPP3R1* | MAPK signaling: phosphatase that binds $Ca^{2+}$ and calmodulin | | ↓ | - | 2°RE |
| *CAMK1G* | MAPK signaling: $Ca^{2+}$ and calmodulin dependent kinase | | ↓ | | 1°RE |
| *EFNB1* | Craniofacial development: receptor ligand associated with deformities | ~ | ↓ |  |  |
| *SOX14* | Craniofacial development: transcription factor associated with deformities |  | ↓ |  | 2°RE |
| *BNC2* | Melanocyte zinc finger protein affecting human skin color saturation | ~ | ↓ |  | 2°RE |

### Divergent peaks in regions of low recombination

Peaks containing 6/31 candidate genes for coastal adaptation coincided with centromeric regions in Zebra Finch (Table S6), and in 4/6 of these regions we detected a pattern in which sub-regions with higher *F*_ST_ and low-to-zero π abruptly alternated with subregions of higher π and low *F*_ST_ (Scaffold 10, Figure 4b; Scaffold 44, Figure S2f; Scaffold 137, Figure S4b; Scaffold 148, Figure 5b). This pattern was present at only one non-centromeric candidate gene peak (Scaffold 127, Figure S3f).

**Figure 8.**
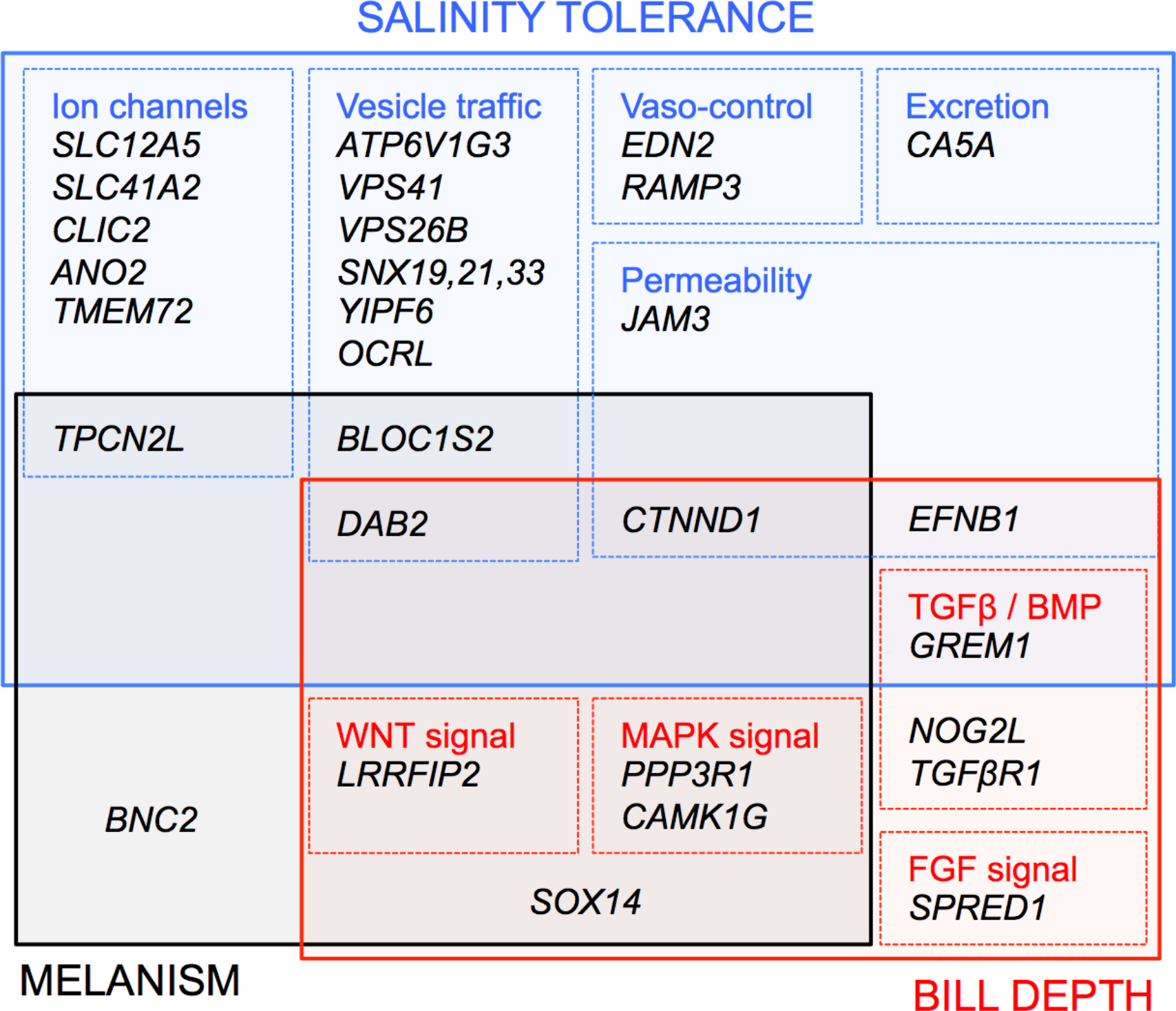
Functional categories encompassing candidate genes for tidal marsh adaptation in swamp sparrows (N=31). In several cases, candidates from one functional category may exert pleiotropic phenotypic effects on other adaptive traits.

## Discussion

### Inferring demography and selection from genome-wide patterns

Genome-wide *F*_ST_ and nucleotide diversity in inland and coastal swamp sparrows was very low, consistent with a very recent split, high levels of incomplete lineage sorting, ongoing hybridization, or all of the above. In coastal birds, on average, Tajima’s D was negative in regions outside of divergence peaks, representing an excess of rare alleles across the genome. This pattern is consistent with a bottleneck having occurred during coastal colonization followed by recent population expansion (Tajima 1989). Although negative genome-wide Tajima’s D provides further support for the demographic context proposed for the evolutionary origin of coastal swamp sparrows (very recent colonization and local adaptation), it might compromise our power to detect regions of negative D due to locus-specific selective sweeps. Runs of homozygosity and nucleotide diversity therefore provided additional lines of evidence with which to identify the signature of locus-specific sweeps. All 31 candidate loci located at *F*_ST_ peaks contained runs of homozygosity, reduced nucleotide diversity, or both.

Several candidate genes that exhibited evidence of a selective sweep via runs of homozygosity and reduced diversity were characterized by positive instead of negative values of Tajima’s D within coastal birds (9/31), a departure from the overall negative genomic background that indicates either balancing selection or structure at that locus within coastal populations (Table 2). Since there is no genome-wide signal of structure due to drift or isolation-by-distance across breeding sites within the coastal population (Deane-Coe et al. 2018), scenarios that could explain this pattern at certain candidate genes include 1) the presence of several different haplotypes in the coastal population that each harbor adaptive variants, or 2) selection for different haplotypes at different coastal breeding sites. We found evidence of allele frequency divergence between coastal breeding sites at three of these loci (ATP6V1G3, SPRED1 and CLIC2), but the remainder (including TGFβR1, BLOC1S2, TMEM72 and RAMP3) exhibited minimal allele frequency divergence within coastal birds. The combined signatures of inland/coastal *F*_ST_ divergence, runs of homozygosity, low diversity and positive Tajima’s D can be explained if adaptive variation at these particular genes was present on more than one haplotype background at the time of coastal colonization, and if each standing variant has since been driven to higher frequency by selection in coastal populations. The biogeographic context of swamp sparrow divergence over longer timescales could have provided a source of that adaptive standing variation on several different genetic backgrounds: coastal tidal marsh habitat has likely established and disappeared in sync with several glacial cycles over time (Malamud-Roam et al. 2006), providing an opportunity for periods of local adaptation followed by reabsorption into the pool of variation present in inland populations during time spent in shared glacial refugia. Genes bearing positive values of Tajima’s D may therefore represent those that have been under selection during previous interglacial episodes of coastal adaptation by swamp sparrows, and persisted as standing variation in the population that recently re-colonized the coast.

### Accounting for alternative processes when interpreting F_ST_ peaks

In this comparison of locally adapted swamp sparrow genomes, we interpret peaks of divergence in the relative measure *F*_ST_ that coincide with other divergence in other metrics (ROH, reduced π, Tajima’s D) as targets of natural selection. However, background purifying selection against deleterious mutations in genes located in centromeres, inversions or other low recombination regions is also capable of purging variation and generating *F*_ST_ peaks across the genome that do not represent regions important for local adaptation (Charlesworth et al. 1993; Charlesworth 1998; Noor and Bennet 2010; Nachman and Payseur 2012; Cruickshank and Hahn 2014). We acknowledge this alternative, but since background selection is often considered an effect of linked selection with weakly deleterious mutations (Nordberg et al. 1996; Charlesworth 1998), and divergence between inland and coastal lineages is so recent, it is unlikely that there has been sufficient time for this gradual process (Wolf and Ellegren 2016). Most divergence peaks harboring candidate genes in our study did not coincide with known centromeric regions in the zebra finch (25/31), but a subset did (TGFβR1, SLC41A2, ANO2, ATP6V1G3, RAMP3, NOG2L). However, even those that occurred in low recombination regions contained long runs of homozygosity localized to the peak (5/6), a fingerprint of selective sweeps that is not readily mimicked by background selection. Peaks at these genes were also small relative to the putative region of reduced recombination around centromeres. These lines of evidence suggest that divergent natural selection, not background purifying selection, is largely responsible for peaks of divergence between inland and coastal swamp sparrow genomes, and genes located at these peaks are therefore robust functional candidates for coastal adaptation. In fact, reduced recombination may have facilitated divergence at the subset of candidate genes located at centromeres (Butlin 2005; Hoffman and Rieseberg 2008; Ellegren 2012), particularly since inland and coastal swamp sparrows experience active contemporary gene flow at the ecotone between freshwater and brackish marshes (Turner et al. 2005; Noor and Bennett 2009).

### BMP signaling and bill depth, a putative magic trait

Three central participants in the TGFβ / BMP signaling cascade emerged as candidates in our genome scan: TGFβ receptor 1 (TGFβR1)and two binding antagonists of BMP4 (NOG2L, GREM1). TGFβR1 propagates the BMP signaling cascade by phosphorylating SMAD2 (Von Bubnoff and Cho 2001). Noggin (NOG) is an antagonist inhibitor that binds BMP4 (McMahon et al. 1998; Groppe et al. 2002) and diffuses readily through membranes to create morphogenic gradients in BMP4 signal during development (Jones and Smith 1998). Over-expression of NOG causes birds to grow smaller, thinner bills (Abzhanov et al. 2004). The candidate gene under a peak in our genome scan is a Noggin-2 like sequence (NOG2L). Gremlin 1 (GREM1) is another binding antagonist of BMP4, and mutations in GREM1 are strongly associated with human craniofacial defects like a cleft lip or palate (Mostowska et al. 2015; Ludwig et al. 2016). All three are strong candidate genes for bill depth in swamp sparrows since studies in several other birds, including Darwin’s finches, have already established a strong functional connection between this pathway and increases in the depth axis of bills during bone morphogenesis (Abzhanov et al. 2004; Wu et al. 2004, 2006; Brugmann et al. 2010).

The location of divergence peaks at the two antagonists NOG2L and GREM1 suggests selection on downstream regulatory elements that are distal to the promoter, like shadow enhancers. This type of distal secondary enhancer is fairly common (eg. Markstein et al. 2002; Zeitlinger et al. 2007) and often shares redundancy with the primary enhancer. That redundancy makes shadow enhancers more evolutionarily labile than primary enhancers, free to evolve new binding sites or other features that change expression of the target gene (Hong et al. 2008). The location of the divergence peak at TGFβR1 suggests selection on a coding region, influencing the structure and function of the receptor. Divergence at secondary regulatory elements of BMP antagonists, and at coding regions of the TGFβ receptor, may therefore represent the molecular mechanisms by which natural selection is driving adaptive bill divergence in coastal swamp sparrows. Since selection for deeper coastal bills drives song divergence and positive assortative mating between coastal and inland populations, these molecular changes represent a mechanism by which natural selection could drive ecological speciation between inland and coastal swamp sparrows over evolutionary time.

### A shared mechanism for sparrow plumage and human skin color

The candidate gene BNC2 from scaffold 81 is a transcription factor that acts specifically within melanocytes, and BNC2 is one of the main genetic markers associated with skin color saturation in human populations (Jacobs et al. 2013). In human melanocytes, the allelic variant present at an intergenic SNP proximal to the BNC2 enhancer region determines chromatin accessibility at the enhancer, leading to variation in BNC2 expression in the melanocyte and resulting variation in skin saturation (Visser et al. 2014).

This gene is made even more compelling as a candidate for plumage melanism in coastal swamp sparrows by the observation that coastal birds exhibit increased melanism in their soft parts as well as their plumage, including the skin of their legs and their lower bill (Figure S5). Coastal swamp sparrows experiencing ecological selection for increased melanism may therefore have converged on the same molecular mechanism as human populations. In humans, functional SNPs that associate with skin color fall in an upstream sequence motif that structurally influences chromatin accessibility at the enhancer of BNC2 (Visser et al. 2014). In swamp sparrows we detect a peak of divergence immediately downstream of the gene body, so different molecular features governing BNC2 expression are likely under selection.

### Multiple physiological mechanisms for salt tolerance

Genes with a potential to confer physiological tolerance to salinity constituted the majority of candidates from our genome scan (74%), and these encoded ion transporters, hormones governing kidney vaso-control, regulators of vesicle trafficking, an enzyme affecting water balance during excretion, and adhesion proteins that reduce intestinal permeability to ions. Salt tolerance genes involved in vesicle trafficking were particularly well represented in the list of candidates, and these eight genes are associated with a range of different trafficking networks, vesicle types, transport destinations and vesicle life cycle timepoints.

Two different divergence peaks implicate endothelin and adrenomedullin hormones as being important for coastal adaptation in swamp sparrows due to their effects on kidney nephron vaso-control. Fish express endothelin receptors in their gills and kidneys that modify blood pressure to alter sodium excretion and confer homeostasis (Hyndman 2015). Since changes to blood flow are temporary and reversible, endothelins are particularly important for maintaining osmotic balance in estuarine fish like killifish, where salinity changes dramatically with the tide cycle (Hyndman and Evans 2007, 2009). In rats fed a high salt diet, adrenomedullin was upregulated in the kidney (Cao et al. 2003), conferred protection against kidney damage (Nishikimi et al. 2002) and inhibited a rat’s appetite for salt (Samson and Murphy 1997), indicating that this hormone can regulate osmotic balance by modifying both excretory physiology and behavior. This behavioral effect of adrenomedullin matches well with known mechanisms of salt tolerance in salt marsh specialist sparrows, since they actively decrease drinking rate as salinity increases (Bartholemew and Cade 1963). The peak encompassing RAMP3, a required activator of the adrenomedullin receptor (McLatchie et al. 1998), is the largest in our comparison between inland and coastal swamp sparrows, indicating that natural selection acting on adrenomedullin binding has been particularly strong during local adaptation to tidal marshes. Vasoconstriction and vasodilation hormones may therefore provide a powerful and temporally flexible functional mechanism by which coastal swamp sparrows physiologically respond to changing tidal salinity.

### Pleiotropic effects of salt tolerance genes

Candidate gene BLOC1S2 encodes a subunit of the BLOC1 protein that, in mice, is required to produce normal lysosomes in the kidney and normal melanosomes in the skin. BLOC1 knockout mice exhibited impaired kidney function and a pale coat color (Theriault and Hurley 1970; Nguyen et al. 2002; Dell-Angelica 2004). Candidate gene TPCN2 encodes an ion channel expressed in the kidney, and harbors SNPs that determine blond vs. brown hair in humans (Sulem et al. 2008; Zong et al. 2009). The potential for these two candidate genes to exert pleiotropic effects on both salt tolerance and melanism may explain how melanic plumage has evolved multiple times in bird populations adapting to brackish tidal marshes or saltmarshes. As is the case in swamp sparrows, melanic plumage or pelage may confer an adaptive benefit for saltmarsh mice, shrews, voles, rails and snakes, but pleiotropy with salt tolerance mechanisms may strengthen and simplify natural selection for both traits. Since habitat isolation is often the only barrier between saltmarsh taxa and their inland relatives, this could help adaptive divergence proceed, or be maintained, even in the face of ongoing gene flow (Kondrashov and Mina 1986; Via 2001). Present day inland swamp sparrows exhibit variation in plumage melanism (Greenberg and Droege 1990). If functional variants of these genes were segregating as standing variation in the source population that first colonized tidal marsh habitats, darker birds could have been “pre-adapted” to deal better with salinity and may have experienced differential survival. In other species where saltmarsh melanism has evolved but there is no strong evidence for the adaptive value of that melanism, it could be that melanism is in fact a molecular “spandrel”: a selectively neutral, pleiotropic consequence of selection for salt tolerance (Barrett and Hoekstra 2011; Gould and Lewontin 1979). Future studies characterizing tissue-specific gene expression throughout development are required to test whether BLOC1S2 or TPCN2L do exert pleiotropic effects on swamp sparrow physiology as they do on human and mouse physiology.

A reduction in BMP4 activity due to antagonism by the candidate bill depth gene GREM1 is also required for normal kidney development (Michos et al. 2004, 2007). Like the pleiotropic effect of salt tolerance candidates on melanogenesis, the dual role of the BMP4 antagonist GREM1 in bill and kidney function may help explain an additional axis of parallel evolution among salt marsh birds: larger bill size. Under this scenario the pleiotropic effects of GREM1 on kidney development could have strengthened selection, contributing to patterns of parallel evolution across species like swamp, song and savannah sparrows that have colonized tidal marsh habitat and presumably experienced similar combinations of selective pressures. Alternatively, instead of both traits being adaptive, the convergent evolution of larger bills across tidal marsh sparrows could be a non-adaptive byproduct of selection on GREM1 for only one of its roles, either adaptation to heat stress (bill size) or salt stress (kidney development).

### Peaks without candidates

We did not detect candidate genes for salt tolerance, bill depth or melanism at almost half of the divergence peaks we characterized genome-wide (19/41). Since at least eight other traits distinguish inland and coastal swamp sparrows, including degrees of sexual dimorphism, reproductive biology, molt timing, migration, and the corticosterone stress response (summarized in Deane-Coe et al. 2018), these peaks could harbor functional genes for other coastally adaptive traits. Alternatively, they could harbor functional genes for traits under selection in inland freshwater swamp sparrows.

## Conclusions

Swamp sparrows have recently colonized coastal tidal marshes and have rapidly evolved a suite of local adaptations. We detected signatures of selection on 20 different salt tolerance genes representing at least five different physiological mechanisms; a diverse panel of candidates that speaks to the multigenic nature of adaptation to a systemic environmental stressor like salinity. We also detected selection at genes with important evolutionary roles in other systems, including BMP signaling factors that influence bill depth in Darwin’s finches and the melanocyte transcription factor BNC2 that determines human skin tone. The targets of natural selection for swamp sparrow bill depth and plumage melanism are therefore examples of taxonomically broad functional convergence, and are evidence of the repeatability of evolution. The results of our genome scan also emphasize the perhaps-underappreciated role of pleiotropy in local adaptation. Six of the candidate genes for salt tolerance also have well-documented effects on bill shape and plumage melanism. In cases like the swamp sparrow, in which the adaptive value of deep bills or melanism has been tested and demonstrated (Olsen et al. 2008a; Peele et al. 2009; Greenberg et al. 2012), pleiotropy with salt tolerance mechanisms may have facilitated divergence by offering natural selection a single target capable of influencing several adaptive phenotypes. Alternatively, pleiotropy could cause divergence in a suite of traits as a byproduct of selection on a single trait, and this needs to be incorporated as a component of null models in future studies of adaptive phenotypic divergence.

## Acknowledgements

The authors thank L. Campagna, D. Toews, and J. Walsh for comments and discussions that improved this manuscript. The Smithsonian Conservation Biology Institute’s Center for Conservation and Evolutionary Genetics (R. Fleischer), and The Cornell University Museum of Vertebrates (C. Dardia, V. Rohwer) contributed samples. Blackwater National Wildlife Refuge (M. Whitbeck) and Fishing Bay Wildlife Management Area (J. Moulis) provided logistical support, and A. Bessler, K. Bostwick and K. Deane-Coe provided field assistance. John Anderton graciously permitted reuse of his illustrations of inland and coastal swamp sparrows. This research was supported by the Athena Fund at the Laboratory of Ornithology, Cornell Vertebrate Genomics, and Doctoral Dissertation Improvement Grant No. 1501471 from the National Science Foundation. Fellowship support for P.D. was provided by the David and Sandra Junkin / Walter E. Benning Graduate Fellowship at the Cornell Laboratory of Ornithology and the Cornell University Andrew and Margaret Paul Graduate Fellowship. Sampling was conducted under appropriate state and federal permits and all procedures conformed to protocols approved by Cornell’s Institutional Animal Care and Use Committee (IACUC).

